# SAW: An efficient and accurate data analysis workflow for Stereo-seq spatial transcriptomics

**DOI:** 10.1101/2023.08.20.554064

**Authors:** Chun Gong, Shengkang Li, Leying Wang, Fuxiang Zhao, Shuangsang Fang, Dong Yuan, Zijian Zhao, Qiqi He, Mei Li, Weiqing Liu, Zhaoxun Li, Hongqing Xie, Sha Liao, Ao Chen, Yong Zhang, Yuxiang Li, Xun Xu

**Affiliations:** BGI-Shenzhen, Shenzhen, Guangdong, China; BGI-Beijing, Beijing 102601, China; BGI-Wuhan, Wuhan, Hubei, China

## Abstract

The basic analysis steps of spatial transcriptomics involve obtaining gene expression information from both space and cells. This process requires a set of tools to be completed, and existing tools face performance issues when dealing with large data sets. These issues include computationally intensive spatial localization, RNA genome alignment, and excessive memory usage in large chip scenarios. These problems affect the applicability and efficiency of the process. To address these issues, a high-performance and accurate spatial transcriptomics data analysis workflow called Stereo-Seq Analysis Workflow (SAW) has been developed for the Stereo-Seq technology developed by BGI. This workflow includes mRNA spatial position reconstruction, genome alignment, gene expression matrix generation and clustering, and generate results files in a universal format for subsequent personalized analysis. The excutation time for the entire analysis process is ∼148 minutes on 1G reads 1*1 cm chip test data, 1.8 times faster than unoptimized workflow.

## Statement of Need

Stereo-Seq of BGI STOmics^1^ is a panoramic spatial transcriptome technology that achieves ultra-high throughput and ultra-high precision. By capturing mRNA in tissues with the Stereo-seq chip and restoring it to its spatial location, in situ sequencing of tissues is achieved, laying the foundation for a deeper understanding of the relationship between gene expression, morphology, and local environment of cells.

Due to its ultra-high throughput and ultra-high precision, Stereo-Seq generates a large amount of data, which poses a challenge for data analysis. Therefore, efficient analysis tools are needed to solve this problem. In addition, accurate spatial positioning is an important part of data analysis, so accurate positioning will lay a good foundation for subsequent analysis.

In the spatial transcriptomics analysis process, large amounts of data can bring some performance issues to the traditional analysis process. Firstly, the alignment of mRNA sequences, regardless of whether using STAR^2^ or other software, cannot meet the performance requirements in the current situation. In the s1 (1cm*1cm) chip, this step can account for 70% of the process time. In addition, the CID mapping step is also an important step in the process, and its accuracy affects the efficiency of spatial positioning. In this step, the information of CID and coordinates needs to be recorded in memory for real-time query and spatial positioning of reads. Faced with large chips, such as S6 (6cm*6cm), the spatial coordinate points can reach as many as 15 billion, and the data structure that stores the correspondence between CID and coordinates will occupy huge memory. Moreover, querying in such a large table will also be slow, especially when we consider fault tolerance, the computational complexity and time consumption will be further increased. Finally, on large chips, matrix operations of the same size as the chip will also have performance bottlenecks such as excessive memory usage and slow speed. These problems need to be solved through high-performance computing technology.

We developed Stereo-Seq Analysis Workflow (SAW) standard analysis process which takes FASTQ^3,4^ as inputs, and goes through mRNA spatial location restoration, filtering, mRNA genome alignment, gene region annotation, MID (Molecule Identity) correction, expression matrix generation, tissue region extraction, clustering, saturation analysis, and report generation to obtain the gene expression and spatial information of tissues, completing the basic analysis of spatial transcriptomics data.

## Implementation

### Processing and parallelization of spatial information in large chips

The principle of spatial localization of sequencing data by spatial transcriptomics is to mark the spatial position and sequencing reads with a 25bp Coordinate ID (CID) sequence, and then locate the sequencing reads back to their original spatial position by matching the CID sequence on the reads. However, due to the fact that the DNA sequence obtained by current sequencers is not 100% accurate and has a certain error rate, error tolerance is required when matching the CID sequence. The current error tolerance strategy is to replace each base on the CID sequence with the other three bases (the gene sequence is composed of four bases A/G/C/T), and then perform the matching.

Due to the high resolution and large field of view of Stereo-Seq itself, the number of spatial coordinate points is very large. For example, for the S6 chip (6cm*6cm), the number of spatial coordinate points can reach as many as 15 billion. Simply storing the corresponding relationship between each coordinate point and CID sequence in a data structure will consume more than 600G of memory, and the query speed is very slow, seriously affecting the applicability and analysis efficiency of standard analysis.

Therefore, we have split the information of spatial coordinates and CID which stored in mask file, and correspondingly, the FASTQ files are also split according to the same rules. For example, if the mask file need be split into 4 parts, the first base of the CID sequence can be used as the classification criterion and split it into 4 parts starting with A, C, G, and T. If 16 parts was needed, the first two bases can be used as the classification criterion. If a non-power-of-4 number of parts was needed, such as 10 parts, we can use a modulo operation. Similarly, the FASTQ file will be split according to the CID sequence using the same rule, and the corresponding mask file and FASTQ file belonging to the same category will undergo CID mapping. This solves the above memory problem and improves the parallelization of data processing (Figure 1). The Mask file (which records the corresponding relationship between CID sequence and spatial coordinates) and the FASTQ split are paired for subsequent analysis, and then merged when needed.

**Figure 1.**
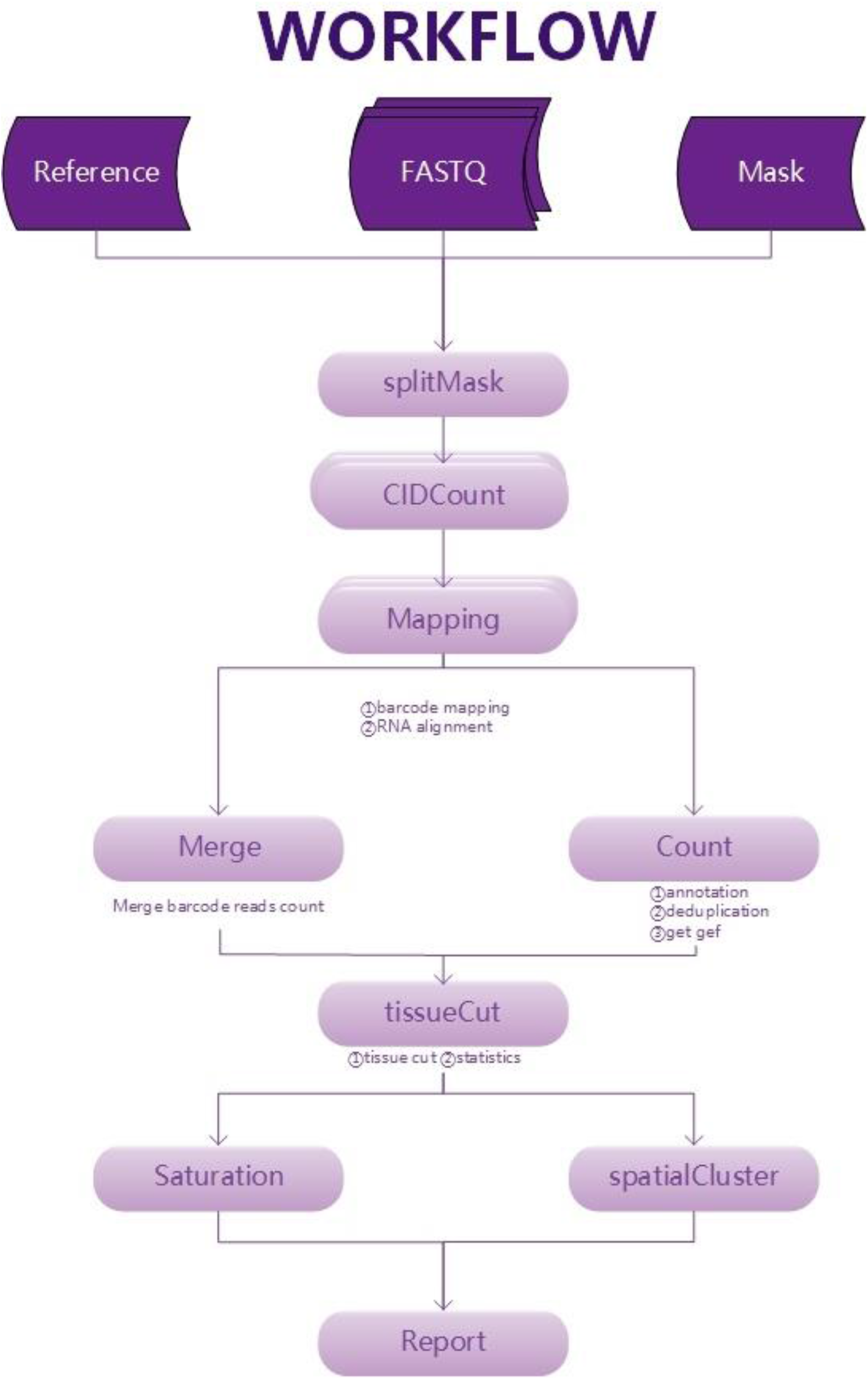
Stereo-seq data analysis workflow

### Rapid alignment of genomes

The successful positioning of spatial location in read will go through several filtering steps, including whether MID contains N bases, whether MID is polyA, MID quality value and whether mRNA contains polyA. Reads filtered through these steps will undergo genome alignment and output a BAM^5^ file containing alignment information. Currently, commonly used RNA alignment tools include STAR, Hisat2^6^, and TopHat2^7^. Among them, STAR is known for its high unique alignment rate and relatively fast speed, but it still cannot meet our requirements when facing spatial omics big data. Therefore, we have made a series of optimization attempts, including using efficient multi-threaded IO models, single instruction multiple data (SIMD), improving L2 cache hit rate and other micro-architectural optimization techniques, redesigning business-level algorithms in data processing, and using FM-index technology in maximum matching prefix search, ultimately accelerating it by 2 times.

### Gene expression matrix generation

Gene expression quantification analysis of STOmics is achieved through the count tool in the analysis software. Count annotates uniquely mapped reads based on mapping alignment results, combined with the reference gene annotation file (GFF/GTF^8,9^) of the corresponding species, and corrects and deduplicates MID, generating processed BAM files, gene annotation, and MID correction and deduplication.

#### Gene region annotation process

1. For each read, search for overlap with the gene interval in the annotation file, calculate the genename/strand on the annotation, and determine whether it belongs to EXONIC/INTRONIC/INTERGENIC, and count the number of each type.
2. Parse the cigar information. Obtain the full length of the read and the starting position and length of each align block.
3. Search for all overlapping genes.
4. Determine which gene to choose. For each gene:
  a. For each align block, calculate with each transcript of the current gene, and obtain multiple exoncnt/introncnt based on the length of overlap with exon/non-exon regions. Take the maximum exoncnt (priority is given to exon>intron).
  b. Accumulate the cnts of each align block to obtain the optimal exoncnt/introncnt. If the exoncnt is greater than or equal to 50% of the read length, mark it as EXONIC. Otherwise, if the introncnt is greater than or equal to 50% of the read length, mark it as INTRONIC. Otherwise, mark it as INTERGENIC.
  c. Choose the most reliable gene from multiple genes.
    i. First, obtain a list of genes with the best annotation results (priority is given to EXONIC>INTRONIC>INTERGENIC).
    ii. From these genes, select the gene with the largest overlap as the annotation result.
    iii. If multiple genes have the same overlap length, randomly select one gene (the selection rule is to choose the gene with the smaller start and end).

### MID correction

Correcting the error mid caused by sequencing errors based on Hamming distance, the process is as follows:

1. Data preparation. A nested map of the form {cidgene: {mid: cnt}} is used to store the number of each cid and gene combination under various mids.
2. Correction.
  a. Set parameters. Minimum number of mid types threshold/tolerance number threshold/mid length, default 5/1/10.
  b. Correction. For each group of data in the nested map:
    i. Check the number of mid types, and continue processing if it is greater than or equal to the threshold.
    ii. Sort in descending order according to the cnt of mid, obtaining a list of the form [(mid1, cnt1), (mid2, cnt2), …].
    iii. Traverse the sorted list in reverse order, starting with the mid with the smallest cnt, and calculate the base error with other mids. If it satisfies the tolerance number threshold, correct the current mid to the mid with a larger cnt and transfer the cnt of the current mid to the correct mid.
    iv. Obtain the nested map after correction in the form of {cidgene: {oldmid: newmid}}.
    Expression matrix
    1. Select reads annotated to EXON or INTRON.
    2. Filter out reads with directions opposite to the annotated gene chain direction.
    3. Group by coordinates, gene, and MID in order.
    4. Count the number of unique MIDs for each coordinate and gene, which is the expression matrix.
3. Example Given a fault tolerance threshold of 1, assuming the sorted MID sequence and count are as follows:

~~~
{
 “AAA”: 5,
 “GGA”: 4,
 “AGA”: 3,
 “AAT”: 2,
 “GGG”: 1,
 “CCC”: 1
}
~~~ Correction process: Calculate the fault tolerance count between “CCC” and “AAA” - “GGG”, all of which are greater than the threshold of 1. Calculate the fault tolerance count between “GGG” and “AAA” - “AAT”, and find that when encountering “GGA”, the fault tolerance count is 1. Then update the count values of both and record the corresponding relationship before and after correction. Calculate the fault tolerance count between “AAT” and “AAA” - “AGA”, and find that when encountering “AAA”, the fault tolerance count is 1. Then update the count values of both and record the corresponding relationship before and after correction. Calculate the fault tolerance count between “AGA” and “AAA” - “GGA”, and find that when encountering “AAA”, the fault tolerance count is 1. Then update the count values of both and record the corresponding relationship before and after correction. Calculate the fault tolerance count between “GGA” and “AAA”, which is greater than the threshold of 1. Update the original data to:

~~~
{
 “AAA”: 10,
 “GGA”: 5,
 “AGA”: 0,
 “AAT”: 0,
 “GGG”: 0,
 “CCC”: 1
}
~~~ Save the mapping relationship before and after correction:

~~~
{
 “AGA”: “AAA”,
 “AAT”: “AAA”,
 “GGG”: “GGA”
}
~~~

### Extracting data of tissue coverage area

Extract the data of tissue coverage area based on the tissue outline. Tissuecut implements two methods of deep learning and traditional image processing, compatible with two types of images: tissue microscopic images and gene expression heat maps, and designs an end-to-end tissue region extraction algorithm. The deep learning method uses the BiSeNet^10,11^ network algorithm of the neural network algorithm to train two lightweight real-time segmentation network models, which are used to extract tissue regions in microscopic images and gene expression heat maps respectively; the traditional image processing algorithm mainly extracts tissue regions based on the grayscale value information of the image. The algorithm process is as follows:

For different types of images to be extracted and different algorithms selected, different image preprocessing processes are used;

Use deep learning or traditional algorithms to extract tissue regions;

Post-process the algorithm results, filter noise, and obtain the final tissue region. Then extract the tissue outline based on the tissue region, and then obtain the data corresponding to the contour coordinates in the region.

### Clustering

A clustering process for identifying heterogeneity and similarity among cells in tissue regions uses spatial information and gene expression levels. The clustering process involves three steps:

1. Data preprocessing: This step involves filtering, normalization, and standardization of the data. The purpose of filtering is to remove cells with too few genes. Normalization and standardization aim to transform the data into the same scale and eliminate the adverse effects of outliers.
2. Feature selection: Principal component analysis (PCA) and umap^12^ are used for feature selection and dimensionality reduction. The most representative genes are selected from all gene expression values for subsequent clustering analysis.
3. Clustering analysis: The unsupervised clustering is performed using the leiden^13^ algorithm, which is a graph-based clustering method. First, a neighborhood graph is constructed based on the similarity between cells, where each cell is considered a node and the connections between nodes represent their similarity. Each node is initially considered as a separate cluster, and the modularity of the entire graph is calculated. Then, adjacent nodes are iteratively merged to improve modularity. In each iteration, the modularity of merging each node with its neighboring nodes is calculated, and the merging method with the highest modularity is selected. After the iterative process converges, the final clustering result is obtained.

## Saturation analysis

### Preparation

The formula for calculating saturation value is 1-(uniq reads/total reads). Sample 5% of bin200 unique coordinates, restore them to bin1 coordinates, and use the sampled bin1 coordinates under tissue to filter data, construct a list of (x, y, gene, mid), and accumulate all count values to obtain anno reads.

### Saturation calculation

Shuffle the previous list, process the data in order according to the sampling interval of {0.05, 0.1, 0.2, 0.3, 0.4, 0.5, 0.6, 0.7, 0.8, 0.9, 1.0}, and for each sampling point, calculate the total reads/saturation value/median number of genes under bin1/bin200 respectively, and output the statistical result file.

#### Saturation value

Calculate the uniq reads (i.e., gene and mid are both uniq) and total reads under all bins using the formula 1-(uniq reads/total reads); the algorithm for bin1 and bin200 is the same.

#### Median number of genes

Calculate the number of uniq genes under each bin and take the median; the algorithm for bin1 and bin200 is the same.

#### uniq reads

The uniq reads of bin1 are all the uniq reads (i.e., gene and mid are both uniq) under all bin1; the uniq reads of bin200 are all the uniq reads under all bin200 (x, y coordinates of bin1, gene, and mid are all uniq).

### Memory issues in large chip scenarios

In large chip scenarios, in addition to the Mask file, storing gene expression information in large file IO and matrix calculations can cause excessive memory consumption. To solve this problem, we have used a series of optimization techniques, including batch processing of large file IO and partial matrix calculations, pre-calculating sizes to avoid using dynamically expanding data structures, and designing more finely tuned custom data types with smaller memory overhead based on business characteristics. These techniques have enabled large chip data to be successfully completed on ordinary memory machines (256G).

## Examples

### mRNA spatial position restoration, filtering, and genome alignment statistics

Taking SS200000135TLD1 data (https://github.com/BGIResearch/SAW/tree/main/testdata) as an example, execute mRNA spatial position restoration, filtering, and genome alignment in sequence, and obtain the statistics shown in Figure 2. Among 1G reads, 818M (78.8%, compared to the previous step) can be aligned back to spatial positions. After filtering, there are still 763M (93.3%) reads left. After alignment to the genome, 641M (84.0%) uniquely mapped reads are obtained.

**Figure 2.**
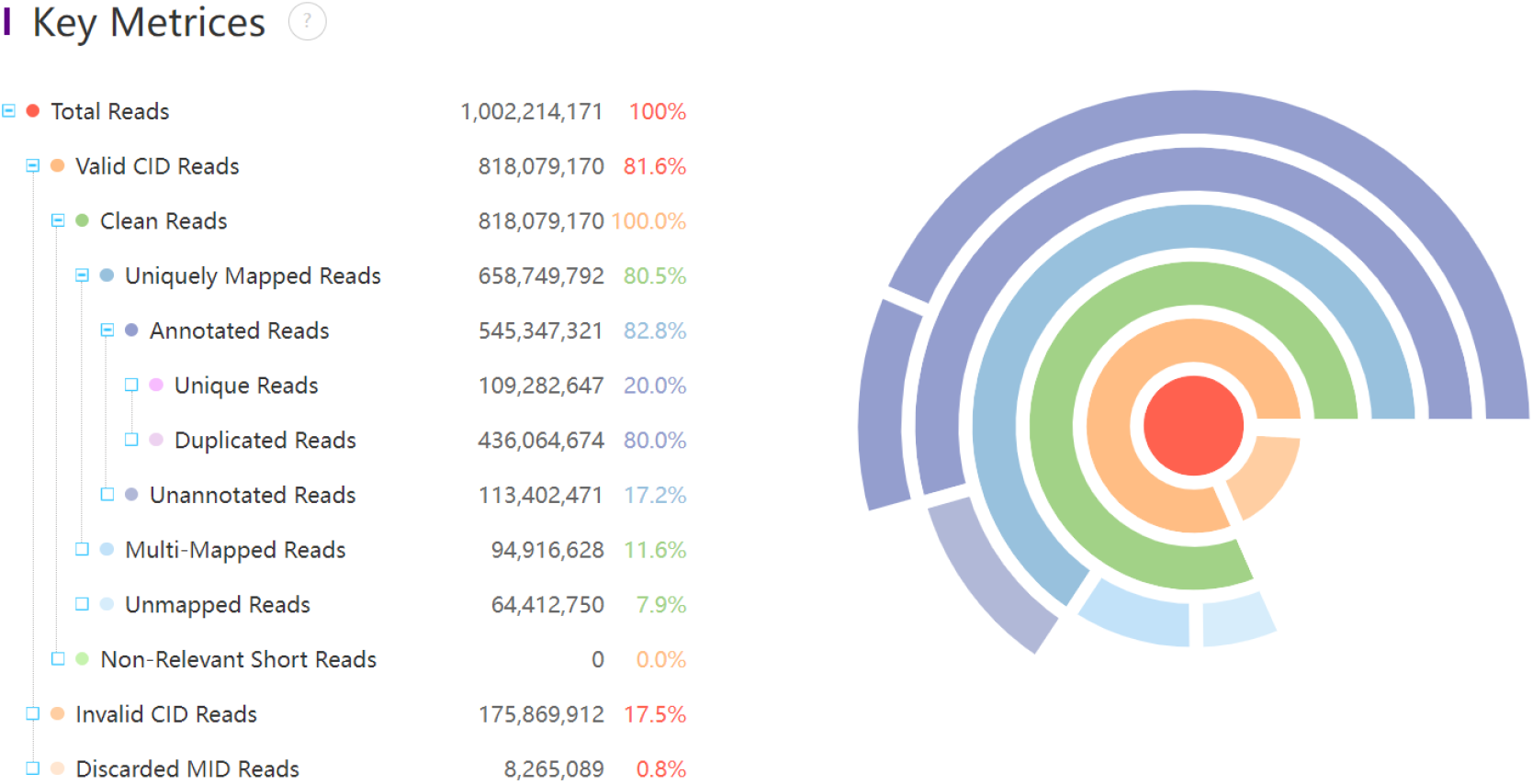
Summary of spatial position restoration, filtering, and genome alignment

### Gene expression spatial distribution map

The bam file generated after genome alignment is annotated and MID-corrected to produce expression information and statistical results. The expression information is stored in hdf5 format and can be visualized (Figure 3). The statistical results provide the number of exonic regions, introns, and intergenic regions annotated. Finally, 481M reads were annotated in the exonic region (Table 1)ο

**Table 1.** A demo of reads annotation statistics.

| type | number |
| --- | --- |
| Exonic | 495,050,694 |
| Intronic | 50,296,627 |
| Intergenic | 113,402,471 |
| Transcriptome | 545,347,321 |
| Antisense | 112,611,349 |

**Figure 3.**
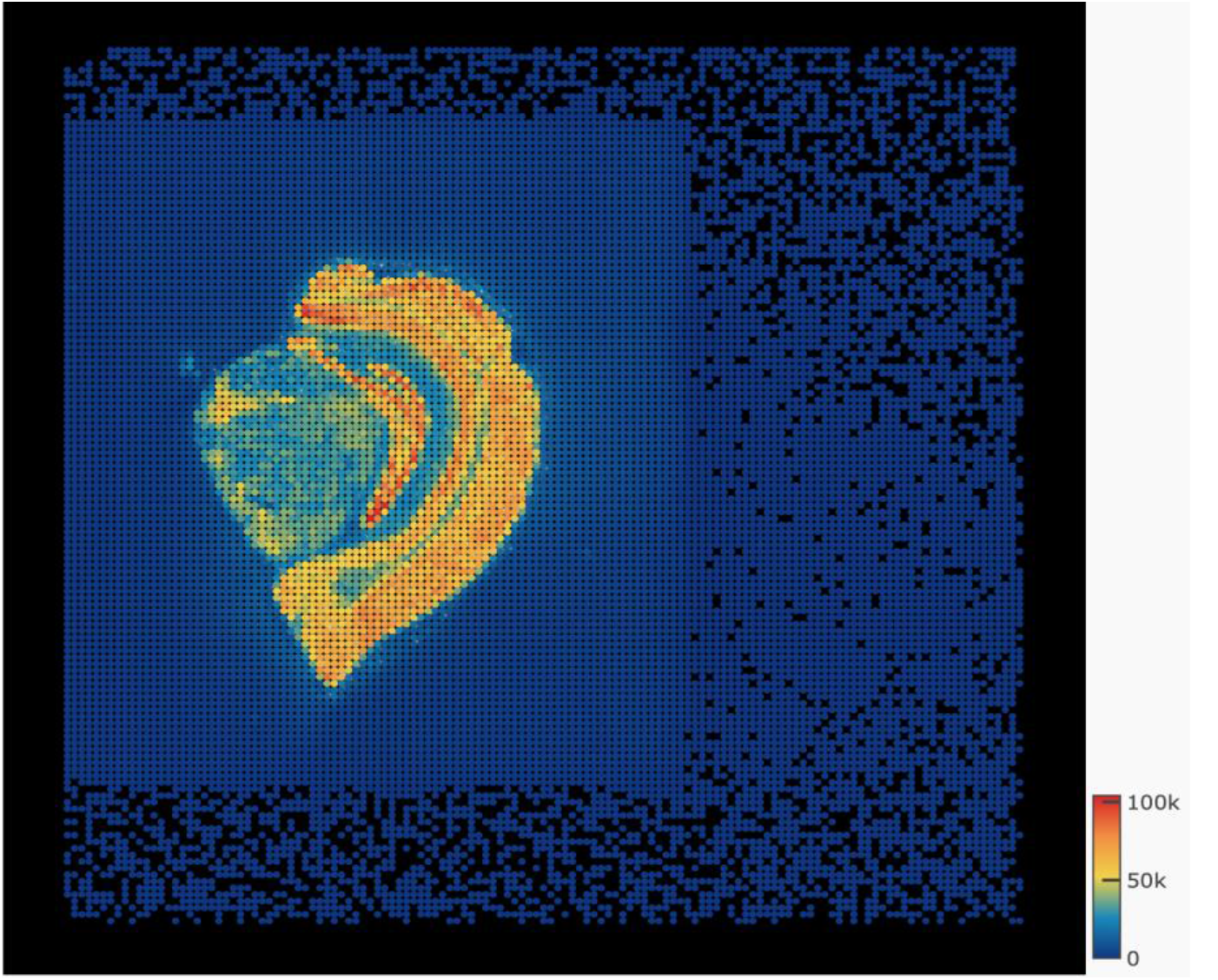
A demo of spatial visualization of gene expression information

### Spatial Clustering

Gene expression profiles are calculated for each position within bin200 (a 200×200 grid of points), and then spatial clustering is performed (Figure 4). This results in 21 classifications, which roughly align with the cell clustering results.

**Figure 4.**
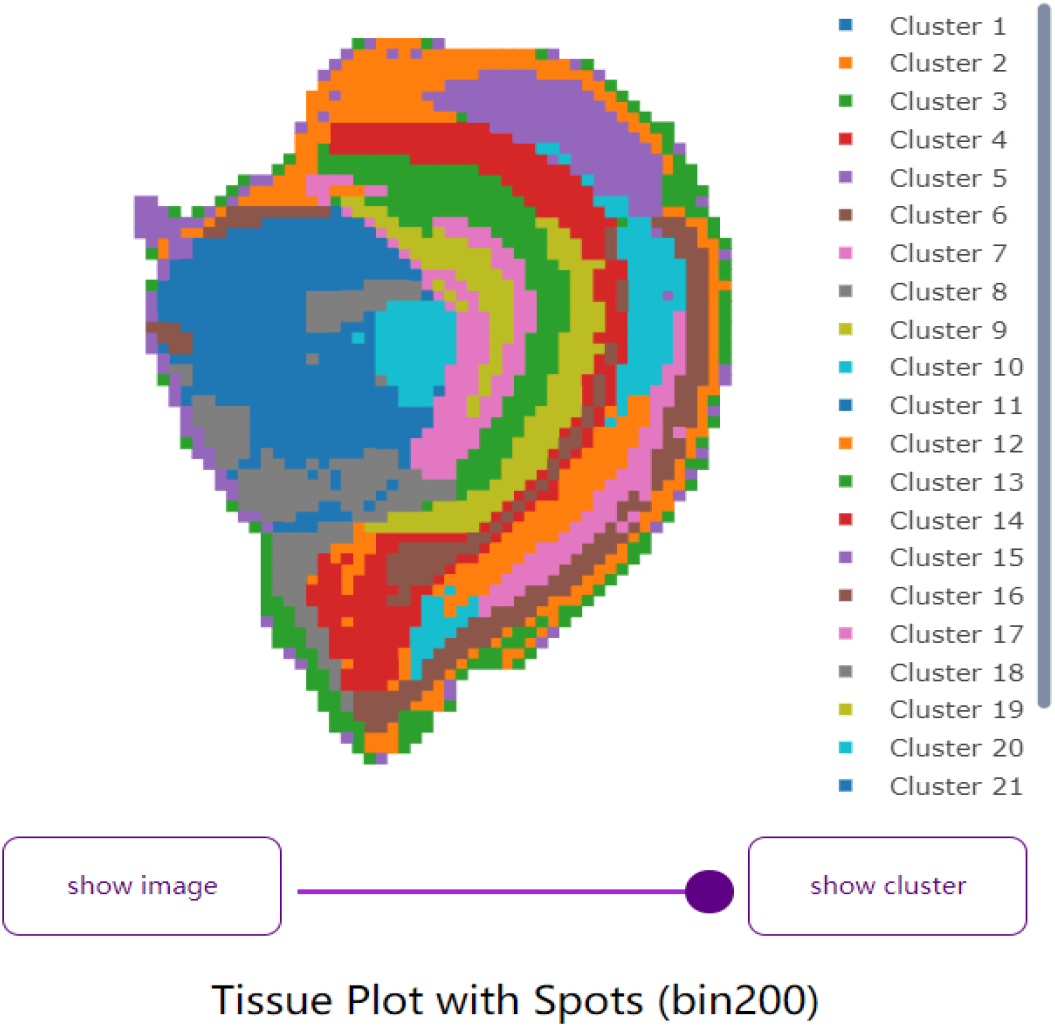
A demo of spatial clustering for mouse brain data

### Saturation Analysis

Saturation analysis shows that the median sequencing depth and number of gene species tend to saturate, while the number of unique reads has not yet saturated (Figure 5).

**Figure 5.**
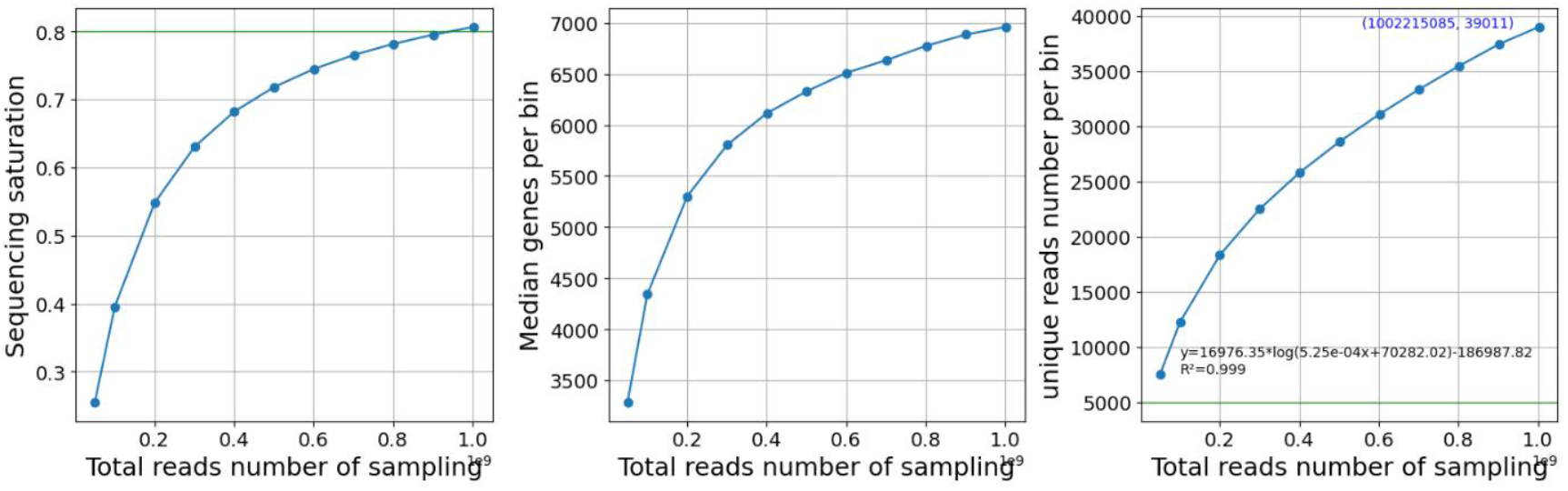
A demo of reads saturation statistics

### Testing

Through a series of high-performance computing techniques, each tool in the pipeline has been optimized. We conducted performance tests on three samples to evaluate changes in runtime and memory. Data1 and data2 are both s1 chips (1cm 1cm) with around 1 billion reads, while data3 is a large chip (2cm*3cm). After optimization, the runtime on data1 decreased from 263.1 minutes to 148.1 minutes, resulting in a 1.8x speed increase. And the time of mapping decreased from 175.0 minutes to 106.9 minutes. On data2, the runtime decreased from 227.9 minutes to 127.3 minutes, resulting in a 1.8x speed increase. And the time of mapping decreased from 158.0 minutes to 83.7 minutes (Table 2). In terms of memory optimization, after process optimization, the memory peak of tissueCut on data3 decreased from well over 83.5G to 33.5G.

**Table 2.** The elapsed time used by analysis workflow before and after optimization.

| performance | Data1(s1 chip,~1G reads) |  | Data2(s1 chip,~1G reads) |  |
| --- | --- | --- | --- | --- |
|  | before optimization | after optimization | before optimization | after optimization |
| elapsed time(min) | 263.1 | 148.1 | 227.9 | 127.3 |

### Future directions

The alignment rate of CID affects the amount of data that enters subsequent analysis. In the test sample, we obtained a successful alignment rate of 78.8% of reads. Due to sequencing errors and alignment algorithm limitations, approximately 20% of reads could not be aligned. In the future, more accurate algorithms (such as those that consider mask CID and fastq CID mismatch base quality values and spatial features) or deep learning models may further improve the accuracy of the pipeline.

## Availability of source code and requirements

Project name: SAW

Project home page:

https://github.com/BGIResearch/saw_tools (tools source code)

https://github.com/STOmics/SAW (script and docker)

Operating system(s): Linux

Programming language: c++, Python

Other requirements: Python >=3.8

License: GNU General Public License version 3

## Data Availability

https://github.com/BGIResearch/SAW/tree/main/test_data

The data that support the findings of this study have been deposited into CNGB Sequence Archive (CNSA)^14^ of China National GeneBank DataBase (CNGBdb)^15^ with accession number CNP0004437.

## Acknowledgements

The authors would like to acknowledge STOmic Cloud(https://cloud.stomics.tech) for supplying software analysis, China National GeneBank and National Key R&D Program of China (2022YFC3400400).

